# Morphological variations in external genitalia do not explain the interspecific reproductive isolation in *Nasonia* species complex (Hymenoptera: Pteromalidae)

**DOI:** 10.1101/2025.10.31.685729

**Authors:** Babita Rahul Baisla, Taruna Verma, Anjali Rana, Ruchira Sen, Rhitoban Raychoudhury

**Affiliations:** Indian Institute of Science Education and Research (IISER Mohali), Knowledge City, Sector 81, SAS Nagar, Mohali, Punjab, India 140306; PG Department of Zoology, Sri Guru Gobind Singh College. Sector 26, Chandigarh

**Keywords:** Aedeagus, aedeagal aperture, sperm length, *N. vitripennis*, *N. longicornis*, *N. giraulti*, *N. Oneida*, female acceptance, prezygotic barrier

## Abstract

External genitalia play a crucial role in the successful copulation of conspecific partners in insects. Due to their structural specificity, external genitalia have often been used for distinguishing closely related species in different insects. However, in the young Pteromalid species complex *Nasonia*, these structures have not received much attention. *Nasonia* is a genus of parasitoid wasps, that lays eggs and completes its life cycle in dipteran pupae. All behavioral studies (including this study) conducted on this genus indicate that varied prezygotic isolation operates among these recently diverged (∼1 million years) species. Whether such prezygotic isolation is caused by the incompatibility in their reproductive organs is not known. Here, we studied the internal male reproductive system of the four species and show that all four species have a similar organisation and thus ruling it out as a potential barrier. Furthermore, to investigate the role of male external genitalia as a prezygotic barrier, we also examined the ultrastructural details, including aedeagal length, aedeagal aperture length, and digital spines. We show that all these parameters and sperm length show variations across the four species. However, these variations do not correlate with the female acceptance of the heterospecific males. Therefore, we conclude that there must be other olfactory or behavioral isolating mechanisms and spatial or temporal factors that maintain the reproductive isolation of these four species.

## Introduction

Interspecific reproductive isolation within a genus can be manifested through morphological, physiological, or behavioral incompatibility between the two potential mates. The morphological features of external genitalia play such a crucial role in the successful mating of insects that their shape and size can be subjected to intense sexual selection (Eberhard, 1985). Moreover, the compatibility and variations in size and shape of external genitalia can also be significant indicators of prezygotic barriers between species. However, often due to the high species diversity, it becomes difficult to study the morphological variations of all species within a genus. Here, we have conducted a comparative analysis of the morphological features of the male external genitalia of the Pteromalid parasitoid wasp, *Nasonia*, to understand their role in interspecific mating compatibility.

*Nasonia* has four reported species: *N. vitripennis, N. longicornis, N. giraulti* (Darling & Werren, 1990), and *N. oneida* (Raychoudhury *et al*., 2010), which diverged from each other within the last 1.0 million years. *N. vitripennis* diverged about 1.0 million years ago, while *N. longicornis* and *N. giraulti* diverged about 0.4 million years ago, and *N. oneida* diverged from *N. giraulti* about 0.35 million years ago (Raychoudhury *et al*., 2010; Werren *et al*., 2010). As a young species complex, *Nasonia* has been extensively used for many studies, including genetic mechanisms of host-parasite interactions (Chabora, 1970; Desjardins *et al*., 2010; Brucker & Bordenstein, 2012), host-parasitoid dynamics (Cornell & Pimentel, 1978; Zareh et al., 1980; Jones & Turner, 1987; Velzen et al., 2016), male behavior (Prazapati *et al*., 2022; Tiwary *et al*., 2022; Verma *et al*., 2025), genomics (Werren *et al*., 2010; Rana *et al*., 2025) and speciation (Breeuwer & Werren, 1990; Campbell *et al*., 1994; Gadau *et al*., 1999; Bordenstein *et al*., 2001; Raychoudhury *et al*., 2010; Werren *et al*., 2010). These closely related species can also be used for studying the roles of external genitalia in prezygotic barriers and speciation. However, very little is known about the variations in the external genitalia of *Nasonia* males. Therefore, we studied the male reproductive system and external genitalia of the four species of *Nasonia*.

The male reproductive system of the flagship species, *N. vitripennis*, consists of a pair of single-lobed testes, vas deferens, seminal vesicles, male accessory glands, ejaculatory duct and the external genitalia or aedeagus (Liu *et al*., 2017). However, whether and how the reproductive system of *N. vitripennis* differs from the other three species is not known. To study the possible variations in external genitalia, we measured the aedeagus length, aedeagal aperture length, and the number of spines on the aedeagal sheaths, along with sperm length. We uncover significant differences across the species in one or more parameters. To understand whether the significant morphological differences pose a prezygotic barrier among the species, we also observed the interspecific and heterospecific mating behavior and tested the correlation between female acceptance with the morphological parameters.

## Methodology

### Strains used

We conducted morphological studies across the males of four *Nasonia* species. The strains used were NV-Asymcx for *N. vitripennis*, NL-IV7 for *N. longicornis*, NG-RV2XU for *N. giraulti*, and NO-NY11/36 of *N. oneida* (Figure 1, upper panel). We maintained all strains at 25°C with 60% relative humidity and at 24-hour constant light. The life cycles of the four species under these conditions were 14 days for *N. vitripennis*, 14.5 days for *N. longicornis*, 15 days for *N. giraulti*, and 16 days for *N. oneida*. We obtained virgin males for the study by hosting <12h old virgin females, on *Sarcophaga dux* (*Diptera: Sarcophagidae*) pupae. To study the external genitalia, we used <12h old virgin males to ensure uniformity and minimise potential variability due to age.

**Figure 1.**
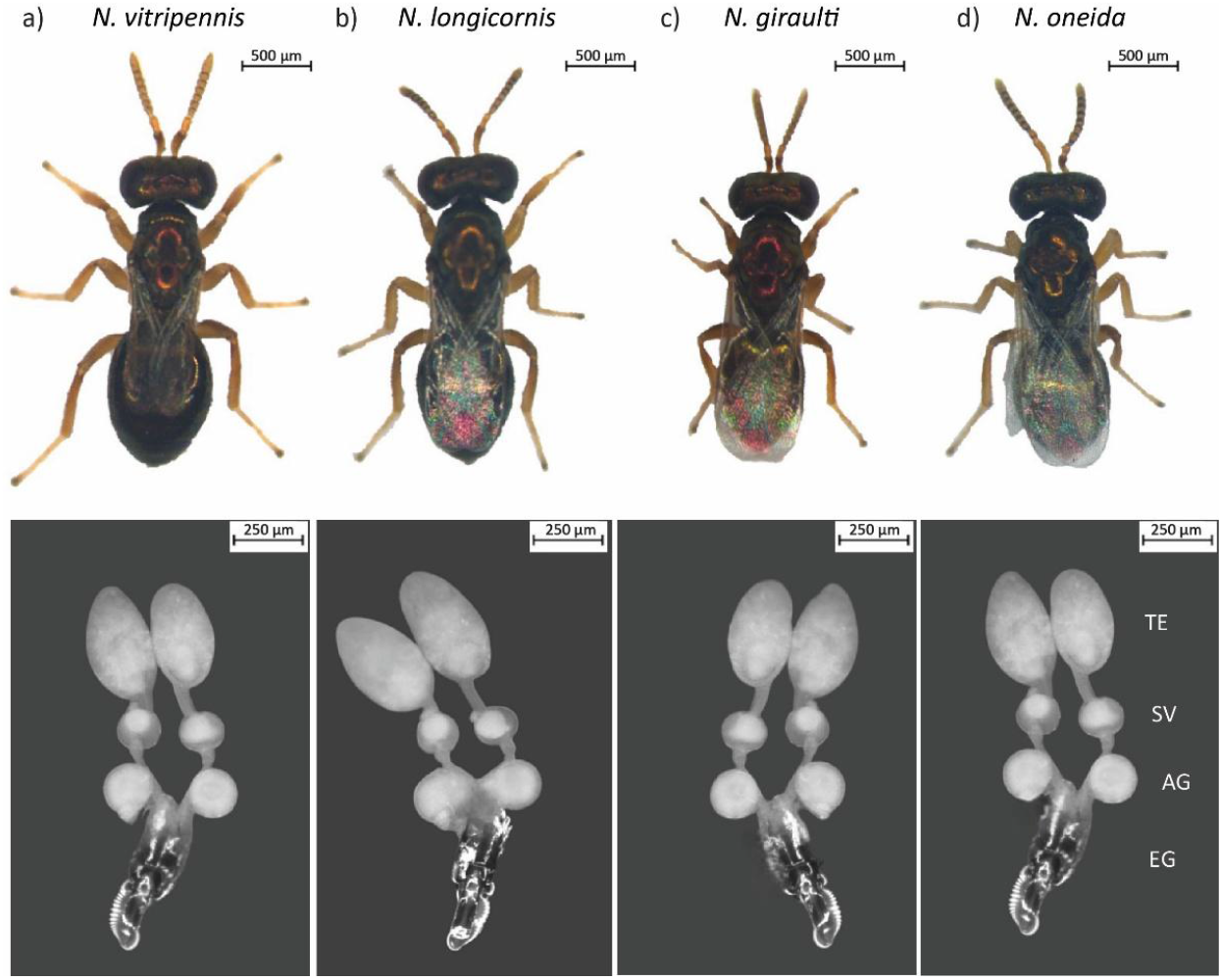
The male reproductive system of the four species of *Nasonia*. The external features of males are represented in the upper panel, and the reproductive system of the corresponding species is presented in the lower panel. TE=Testes, SV= Seminal Vesicle, AG= Accessory Gland, and EG= External Genitalia.

### Male reproductive system and aedeagus length

To study the male reproductive system, we first placed a male in 2µL of 1x Phosphate Buffered Saline (PBS, pH 7.4) and removed the head using a fine needle. The head was then used to measure the interocular distance, which is a reliable estimator of body size (Loehlin *et al*., 2010). We then teased out the male reproductive system from the abdomen using fine needles. We visualised the internal reproductive system and external genitalia through a Leica 10450028 microscope at 50x magnification and measured the aedeagal length at 100x magnification using the Fiji software (https://imagej.net/software/fiji/downloads). The aedeagus lengths were measured from the proximal to distal tip (Figure 2, left panel) and divided by the interocular distance of the same wasp to standardise for the body length variation. The corrected lengths of the aedeagus (n=100 for each species) were then compared statistically across the species.

**Figure 2.**
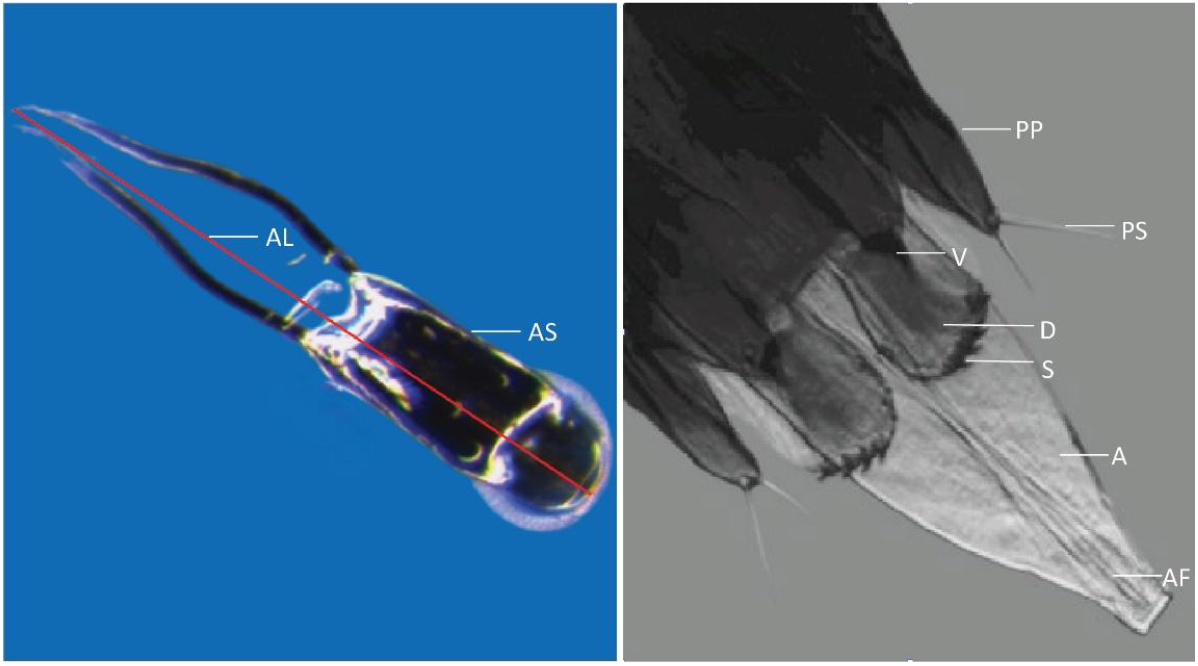
External genitalia within the aedeagal sheath at 50X magnification (left panel) and aedeagus (right panel). AS- Aedeagal sheath, AL- Aedeagus length, PP- Parameral plate, PS- Parameral spine, V- Volsellae, D- Digitus, S- spines, A- Aedeagus and AF- Aedeagal foramen.

### Measurement of aedeagal aperture

We took Scanning Electron Microscopy (SEM) images of the aedeagus for 100 males for each species to measure the aedeagus aperture length. The aedeagus of each male was dissected in 1x PBS under a Leica 10450028 dissection microscope and kept in 2% Paraformaldehyde solution at 4°C. After 12 hours of fixation, they were washed with 1x PBS followed by serial dehydration using 10%, 20%, 30%, 50%, 70%, 90%, and 100% ethanol washes. For maximum dehydration, 2µL Hexa-methyl Disilazane (HMDS) was used after serial dehydration. Aedeagus was then transferred onto carbon tapes and placed on a glass slide. Glass slides were then sputtered (at room temperature for 45 seconds) with gold particles in a gold sputter (Otto Chemie Pvt. Ltd. India) for imaging by SEM. For imaging JOEL Scanning electron microscope (JOEL, JSM 7001F) was used at 12 kV voltage, 10-4 torr vacuum pressure and at different magnifications ranging from 500-3300x at 22°C. The magnification was further adjusted to 1x for standardised measurement.

### Spine count

We used Phase Contrast Microscopy (Olympus), at a magnification of 20x at 22°C, to count the spines present on the digitus (see below). The external genitalia were pulled out and the aedeagal sheath was dissected to visualise the digitus for 50 wasps from each species. The specimens were then incubated for 15 minutes in 1:1 acetic acid and clove oil solution, followed by transfer to a glass slide containing 20μl of Canada Balsam (Otto Chemie Pvt. Ltd. India) to make permanent mounts.

### Staining, visualisation and measurement of sperm

To visualise sperm, we dissected seminal vesicles from males in 1X Phosphate Buffer Saline (PBS, pH 7.4) (PBS). Sperm were released from the seminal vesicles with the help of needles. 20 μl of 2% Paraformaldehyde solution was poured on the released sperm and incubated for 2 hours for fixation. The slides were then washed with 1xPBS solution for 5 minutes to remove the fixative. After washing, they were incubated with 20 μl of 1x PBST (Phosphate Buffer Saline with Tween 20) for 10 minutes. The fixed sperm were washed twice with 1x PBS and stained with 20μl of 20ng/ml of 4′,6-diamidino-2-phenylindole (DAPI) for 30 minutes. After staining, the fixed sperm were washed three times with 1x PBS to remove the excess stain. Then, 20μl of Vectashield (Antifade Mounting Medium, Cole Parmer, United States) was added and stained sperm were covered by a glass coverslip. We stained and measured sperm for 79 *N. vitripennis*, 53 *N. longicornis*, 67 *N. giraulti*, and 58 *N. oneida* males. A Zeiss LSM 980 confocal microscope at a magnification of 400x at 22°C was used for sperm visualisation. For each wasp, the length of one clearly visible sperm was measured twice using the Fiji software (https://imagej.net/software/fiji/downloads), and the average length was used for statistical comparison.

### Estimation of intra- and interspecific mating acceptance

To estimate the percentage of mating within and between species, we placed a virgin male and a virgin female (both <2 days old) in a covered mating chamber made of transparent perspex (15 mm in diameter, 7 mm high, and covered with a 24 mm coverslip). Individual wasps were used only once for each trial, and the pair was observed for 10 minutes using a stereomicroscope at 16x magnification. 70 pairs of wasps were observed for every combination of species. Each mating chamber was washed with 70% alcohol and air-dried before placing the wasps.

### Statistical Analysis

All length measurements were conducted in micrometres using the Fiji software. We analyzed the variation of four species using the Kruskal-Wallis test, and conducted pairwise comparisons between species using the pairwise Dunn’s *post hoc* test. All statistical analyses were performed using the Origin 2022 software (OriginLab Corp). We carried out the Mantel test using R v. 4.2.2 (R Core Team, 2024) for finding any correlation between the female acceptance and aedeagal length or aedeagal aperture length.

## Results

### Description of the male reproductive system

The structural composition of the male reproductive system was strikingly similar in all four species (Figure 1). The system included two single-lobed testes placed ventrally to the digestive system in the abdomen. A short tubular vas deferens originated from each testis and further inflated to form the seminal vesicle, which was visibly filled with compactly packed sperm. A pair of round male accessory glands was present posterior to the seminal vesicles. The vas deferens of both sides joined to form the common ejaculatory duct, which passed through the external genitalia or aedeagus. The aedeagus was further covered by the aedeagal sheath. The two lobes of the testes were oval in shape, non-homogeneous in texture, and approximately similar in size. The testicular lobes were not covered in any sheath.

### Morphology of the male external genitalia

The male external genitalia of *Nasonia* consists of an aedeagus containing the ejaculatory duct, three pairs of sclerites are present on either side of the aedeagus, digitus, volsellae and parameres (gonostyles) positioned laterally (Figure 2, right panel). The cone-shaped aedeagus in *Nasonia* has an uncovered aedeagal aperture or foramen genitale at the end of the ejaculatory duct. The parameres bear a pair of parameral spines and each digitus bears variable numbers of spines at the distal border (see below). (Figure 2, right panel). During copulation sperm passes through the ejaculatory duct and is released through the aedeagal aperture. The aedeagal aperture is elongated, muscular, and lined with cuticle (Figure 3). Two muscular lobes flank the end of the aperture.

**Figure 3.**
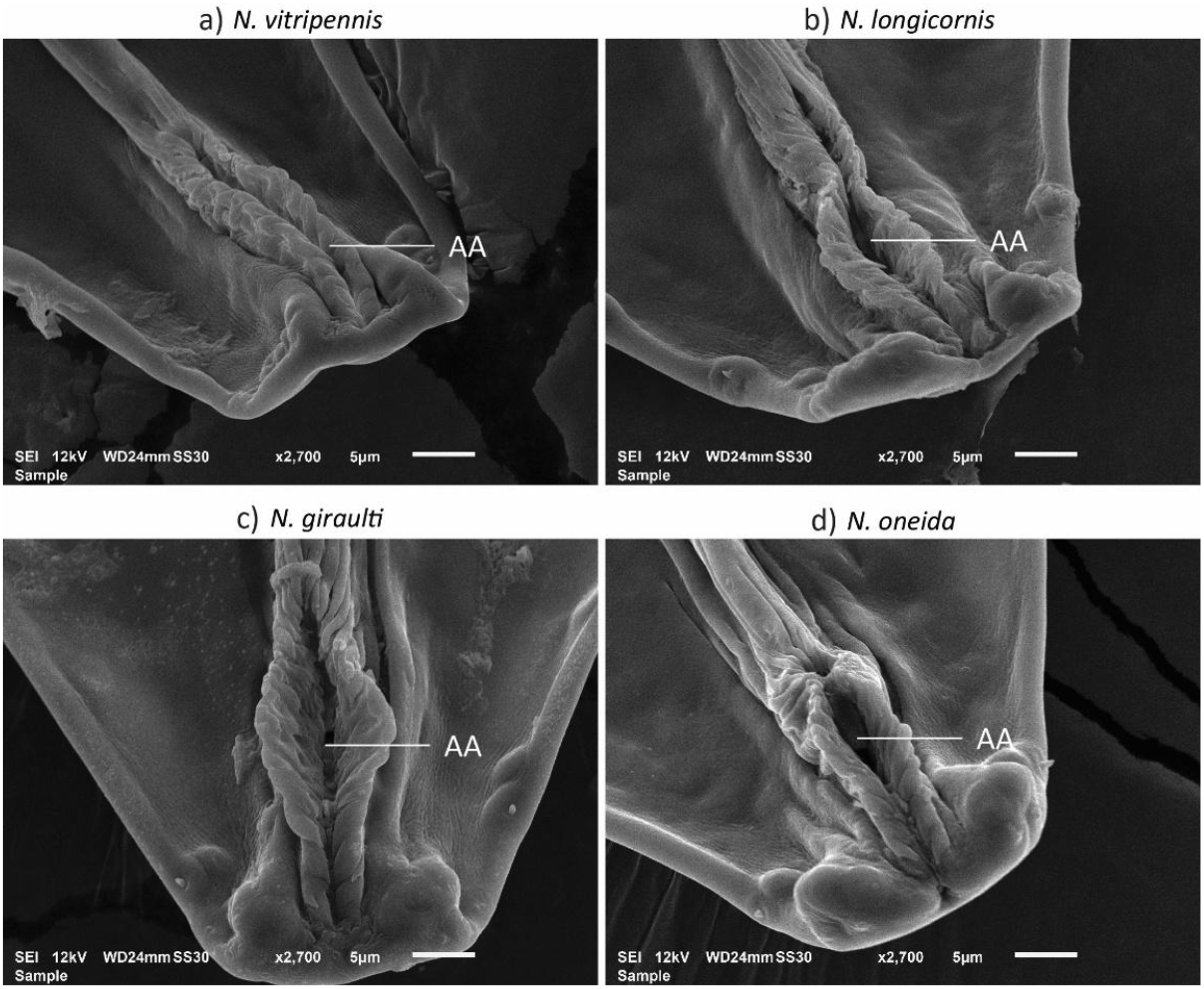
Representative images of the ultrastructural details of the aedeagal aperture in four species of *Nasonia*. AA- Aedeagal aperture.

### Interspecific variation of external genitalia in *Nasonia*

The aedeagus length was longest in *N. longicornis* and shortest in *N. giraulti*, and both were significantly different from *N. vitripennis* and *N. oneida* (Kruskal-Wallis test, *p<0*.*001*; Post-hoc Dunn test, *p<0*.*001* in cases of significant difference, Figure 4a). There was no significant difference in the aedeagus length of *N. vitripennis* and *N. oneida* (Post-hoc Dunn test, *p>0*.*05*).

**Figure 4.**
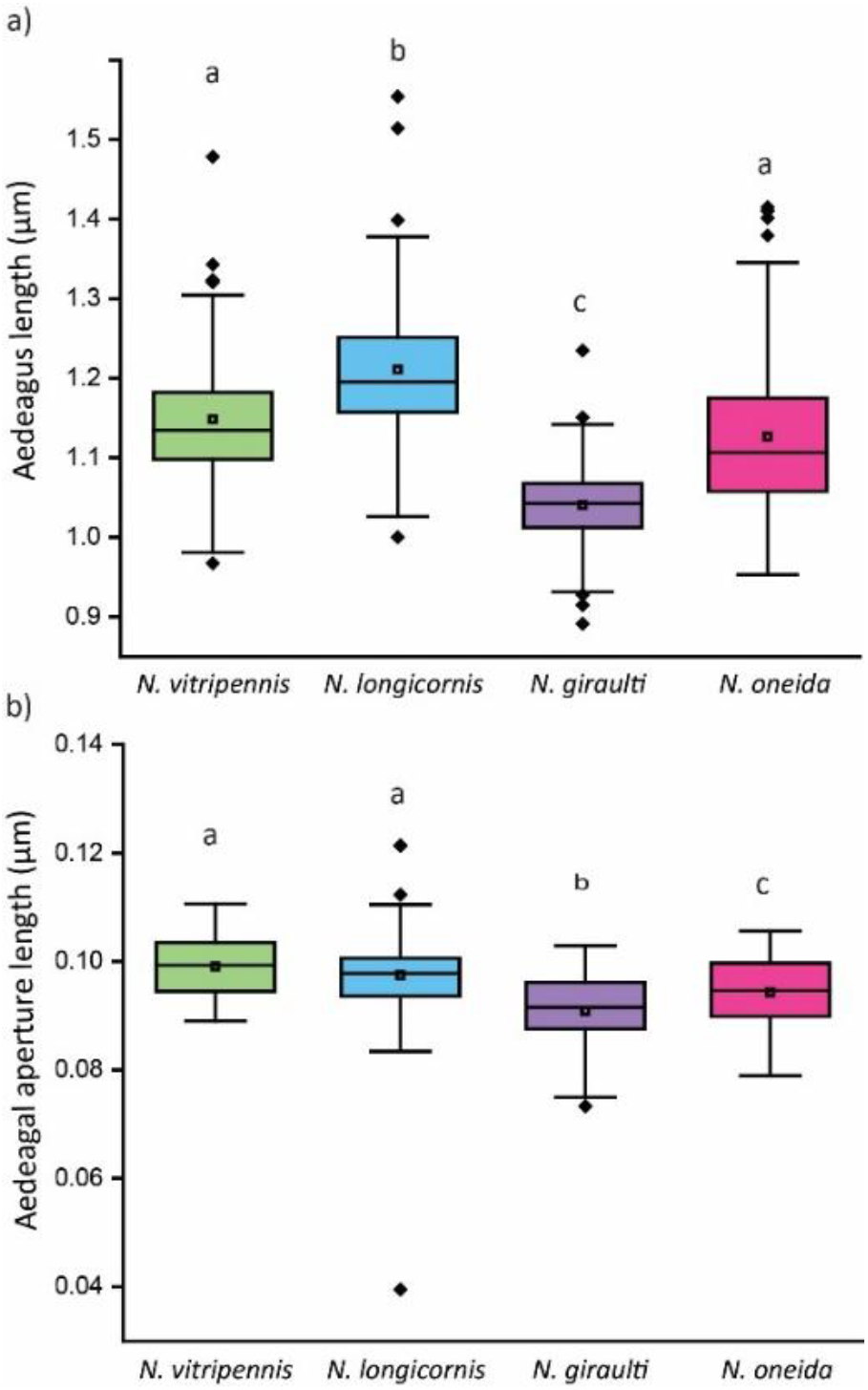
Comparison of (a) aedeagus length and (b) aedeagal aperture length in *Nasonia*. Boxes with different letters are significantly different from each other (Post-hoc Dunn test).

The aperture varied in shape and length within and between species (Kruskal-Wallis test, *p<0*.*001*, Figure 4b). The aedeagus aperture length was not significantly different for *N. vitripennis* and *N. longicornis* (Post-hoc Dunn test, *p>0*.*05*), but both were significantly different from *N. giraulti* and *N. oneida* (Post-hoc Dunn test, *p<0*.*001* for comparisons with *N. vitripennis* and *p<0*.*005* for comparisons with *N. longicornis*). *N. giraulti* and *N. oneida* were also significantly different from each other (*p<0*.*023*).

### Only *N. giraulti* has varying numbers of digital spines

*N. vitripennis, N. longicornis*, and *N. oneida* males consistently had 4 teeth-like spines on each digitus or the sclerite on either side of the aedeagus (Figures 5, upper panel). However, the number of spines varied between 3-5 in *N. giraulti*. Moreover, in some *N. giraulti* wasps, the number of spines also differed between the two sclerites (Figure 5, lower panel).

**Figure 5.**
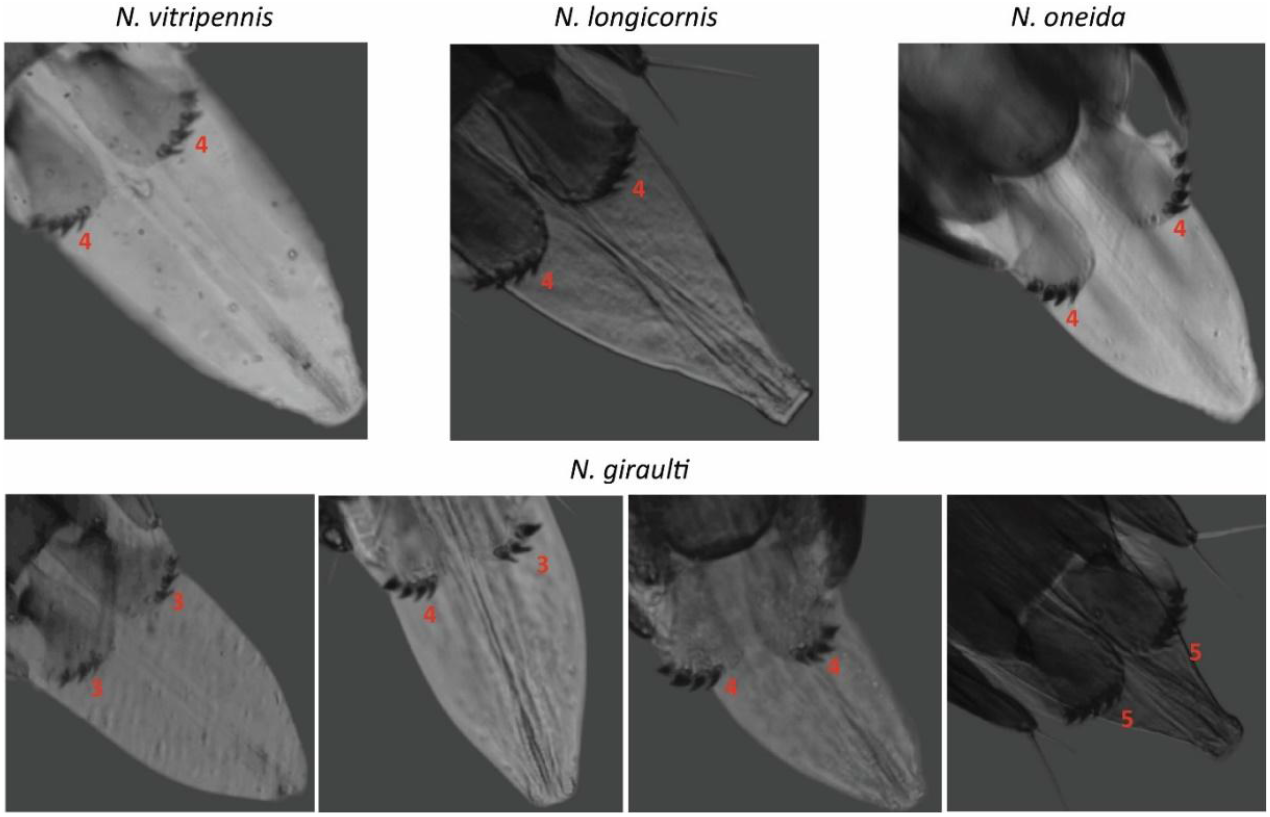
Number of spines (represented by Arabic numbers) on the digitus of *Nasonia*.

### Sperm length - *N. longicornis* has the longest sperm

Sperm lengths varied across the four species of *Nasonia*. The sperm head was slightly thicker with a blunt end, while the tail had a pointy end. *N. longicornis* had the longest sperm, which was significantly different from the sperm length of all other species (Kruskal-Wallis test, *p<0*.*001*, followed by Post-hoc Dunn’s test, Figure 6).

**Figure 6.**
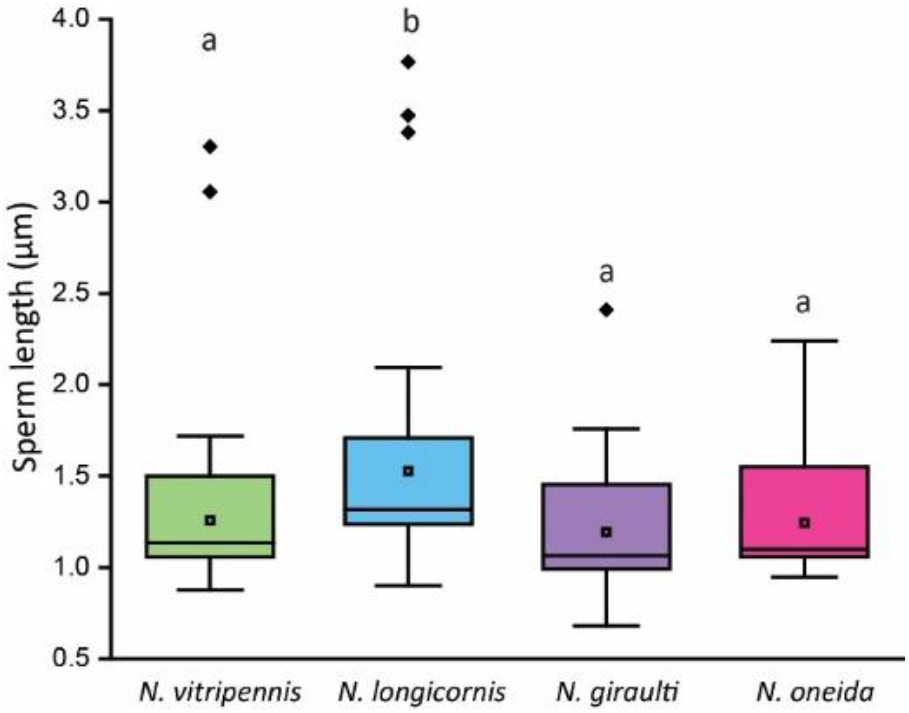
Sperm length variation across *Nasonia*. Boxes with different letters are significantly different from each other (Post-hoc Dunn’s test *p<0*.*001*)

### Female acceptance

In no-choice mating assays, where a virgin male and a virgin female were placed together, complete mating (till the end of post-copulatory behavior) was more frequent in intraspecific (conspecific) pairs. In comparison, the percentage of complete mating in interspecific (heterospecific) pairs varied. *N. longicornis* and *N. giraulti* females showed a greater acceptance of heterospecific males than *N. vitripennis* and *N. oneida* females (Figure 7). Interspecific mating was rarely observed for *N. vitripennis* and *N. oneida* females, as both species were more resistant towards the males of the other species (Figure 7).

**Figure 7.**
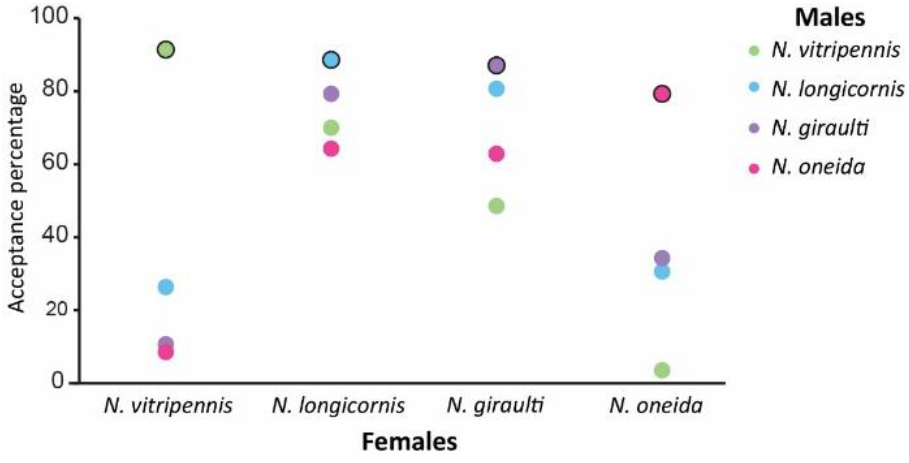
Female acceptance (in percentage) of conspecific males (circles with black borders) and heterospecific males (circles without black borders).

To test whether there was any correlation between the female acceptance and any of the aedeagal parameters, we carried out a Mantel test by making distance matrices using the aedeagal length and aedeagal aperture length. However, there was no significant correlation found between the female acceptance and aedeagal length (r=0.002, *p=0*.*41*), or aedeagal aperture length (r= -0.82, *p=0*.*87*).

## Discussion

The specificity of size and shape of the external genitalia can contribute to maintaining reproductive isolation, sperm transfer, sperm storage, mating success, and paternity(reviewed in Simmons, 2014). Often, the morphology of the external genitalia is so species-specific that they are used to identify and differentiate closely related species (reviewed in Eberhard, 1985; Takami, 2003). We studied the male reproductive system, external genitalia, and sperm length of four species of *Nasonia* to find any differences that may contribute to reproductive isolation. We show that the internal reproductive system of males of the four species is largely similar, with variations in their external genitalia. Previously, Liu *et al* (2017) had documented the male reproductive system of *N. vitripennis*, and we show that, by and large, the same design persists in all four species. This is not surprising as these are a young species group and have diverged relatively recently (Raychoudhury *et al*., 2010). However, in contrast to the photograph published by Liu *et al* (2017), which suggested a spindle-shaped testis and laterally tapered male accessory glands, we found oval-shaped testes and round-shaped male accessory glands in all four species. Our observation suggests that the testes and male accessory glands start to ooze their contents out soon after the dissection, resulting in the distortion of the original shapes. Therefore, for all measuring and documentation purposes, the images should be taken immediately after dissection.

*Nasonia* has a complex ultrastructure of the aedeagus and aedeagal aperture, which were not studied before. However, we assessed these two parameters to investigate if there are any congruent differences in them across the four *Nasonia* species. We found variation in both these parameters across the four species (Figure 4). These data and standardised measuring parameters can be used for further studies, especially in different geographical strains of any one species, to find local variation.

The role of spines on the digitus or the significance of their variation in numbers is currently not known in *Nasonia*. The spines of the external genitalia can play various roles in different insect taxa. In *Anillidris bruchi* males, the digital spines act as a mechano-sensory structure (Cantone & Giulio, 2023). In hymenopteran families like Tiphiidae, Sphecidae, Mutilidae, Scoliidae and Vespidae, spines in genitalia are used for stinging-like behavior by males (Evans & Eberhard, 1970; Eberhard, 1985; Schmidt, 2016) and can also act as anchoring structures to maintain copulatory posture and genital connections (Edvardsson & Tregenza, 2005; Moreno-García & Cordero, 2008; Werner & Simmons, 2008). In several insect species, the digital spines are used to facilitate sperm competition by scooping out the sperm of other males(Hotzy & Arnqvist, 2009) and defend paternity by damaging the female vaginal wall, preventing her from remating (Bergh *et al*.,1992; Miyatake *et al*., 1999). Spines increase the perforation ability of males and the passing of seminal fluid into the female circulatory system, which can increase the chances of paternity and prevent the remating of females (Hotzy *et al*., 2012). Although we are not certain about the roles of digital spines in *Nasonia*, we show that, except for *N. giraulti*, the other three species have four spines on both digitus. Moreover, it is also not clear whether the high rejection of *N. giraulti* males by *N. vitripennis* and *N. oneida* females (Figure 7) was influenced by their variation in spine numbers.

The sperm length varied in the four species, too, but it is unlikely to affect the interspecific mating probabilities. Further study on the ultrastructure of sperm may be useful to determine their exact variation or role, if any, during successful hybridisation.

It is obvious that *Nasonia* females showed a higher acceptance rate for conspecific males and therefore, there are certain prezygotic barriers among the studied strains of the four species of *Nasonia* (Figure 7). Our results were concurrent with the results of previous studies carried out with the same strains (Buellesbach *et al*., 2014) as well as different strains (Giesbers *et al*., 2013). The similar patterns in mating probability among the four species confirm the presence of prezygotic barriers (Raychoudhury, 2015). The earliest divergence of *N. vitripennis* is also reflected in the low acceptance of other males by *N. vitripennis* females, but the females of the most recently diverged species *N. oneida* are equally selective. The within-host mating of *N. giraulti* (Drapeau & Werren, 1999) and the associated non-requirement of selection on females to be discriminating can be a reason why *N. giraulti* females do not discriminate against the males of other species (Drapeau & Werren, 1999; Bordenstein *et al*., 2001; Trienens *et al*., 2021). The study of different parameters here failed to single out any morphological reproductive barrier among the four species of *Nasonia*. However, the significant rejection of heterospecific males by *N. vitripennis* and *N. oneida* females show that the prezygotic barriers, even in a young species complex, can be multifaceted and must have additional behavioral and olfactory factors that influence the decision-making of females.

## Author contribution

BRB and RR conceived and designed the study. BRB collected data, TV and BRB analyzed data and made the figures under RR and RS’s supervision, AR helped with staining of sperm and measuring of sperm length. RS wrote the first draft of the paper with BRB and TV. RS and RR reviewed and edited the manuscript.

## Acknowledgement

The authors thank Prof Sudip Mandal and the central lab facility of IISER Mohali for providing the resources for the SEM.

